# Tumor Treating Fields inhibit cell proliferation and promote immunogenic cell death in biliary tract cancer cells

**DOI:** 10.1101/2025.02.18.638924

**Authors:** Ying Yue, Yingying Wang, Min Yao, Yuanzhen Suo

**Author notes:** Correspondence to: Yuanzhen Suo.

## Abstract

Tumor Treating Fields (TTFields) are low-intensity, intermediate-frequency alternating electric fields that exert antimitotic effects on cancer cells. This study evaluated the in vitro efficacy of TTFields on biliary tract cancer (BTC) cell lines HCCC-9810 and RBE, investigating their sensitivity to TTFields across varying frequencies and electric field intensities. The results demonstrated that a frequency of 150 kHz induced the most pronounced cytotoxic effects, significantly impairing clonogenicity and migratory capacity while inducing abnormal mitotic processes. Furthermore, TTFields treatment triggered a marked upregulation of immunogenic cell death (ICD) biomarkers, including enhanced surface exposure of calreticulin (CRT) and increased extracellular release of high mobility group box 1 (HMGB1) and ATP. These findings indicate that TTFields represent a promising therapeutic approach for BTC, not only by suppressing tumor proliferation but also by enhancing ICD.

## INTRODUCTION

Biliary tract cancer (BTC), encompassing intrahepatic, perihilar, and distal cholangiocarcinoma as well as gallbladder cancer, represents a heterogeneous group of malignancies with limited treatment options and a generally poor prognosis^[1]^. The incidence and mortality rates of BTC are rising globally, posing significant challenges in its clinical management^[2,3]^. Standard therapeutic approaches, such as surgical resection, chemotherapy, and radiation therapy, are often inadequate due to the aggressive nature of the disease, the anatomical complexity of BTC, and the compromised performance status of patients^[4,5]^. In recent years, targeted therapies and immunotherapies have provided new strategies for treating biliary tract cancers^[6,7]^. The TOPAZ-1 clinical trial (NCT03875235) represents a significant advancement in immunotherapy^[8]^. This Phase III study assessed the efficacy of combining durvalumab, an anti-PD-L1 antibody, with the standard gemcitabine and cisplatin (GC) regimen. The trial demonstrated improved overall and progression-free survival (PFS) in patients with BTC who received the durvalumab and GC combination therapy, leading to its approval as a first-line treatment for advanced BTC. However, the efficacy of these treatments remains limited by tumor resistance mechanisms and the immunosuppressive tumor microenvironment.

Tumor Treating Fields (TTFields) represent a novel non-invasive cancer treatment modality that inhibits tumor growth by applying low-intensity, intermediate-frequency alternating electric fields directly to the tumor site. These electric fields are administered to the affected area through transducer arrays positioned on the skin, thereby delivering a localized treatment effect^[9,10]^. The anticancer effects of TTFields primarily stem from the disruption of mitotic spindle formation during cell division, which results in abnormal chromosome segregation and various types of cell death^[11,12]^. The therapy has received FDA approval for treating newly diagnosed and recurrent glioblastoma (GBM), malignant pleural mesothelioma, and as second-line treatment for non-small cell lung cancer (NSCLC) with clinical trial data confirming its efficacy and safety^[13-15]^. In addition to these approved uses, TTFields are currently being studied for a variety of other solid tumors, including pancreatic cancer, ovarian cancer, brain metastases from NSCLC, first-line treatment of NSCLC, and hepatocellular carcinoma (HCC)^[16-20]^. These studies aim to broaden the potential applications of TTFields by evaluating their efficacy when combined with standard of care (SOC).

Immune checkpoint inhibitors (ICIs) have transformed cancer treatment by enhancing immune responses against tumor cells^[21]^. In BTC, ICIs targeting the PD-1/PD-L1 pathway show promise but remain limited by the cancer’s immunosuppressive microenvironment^[22,23]^. Combining TTFields with ICIs represents a promising therapeutic strategy, as TTFields induce immunogenic cell death (ICD), releasing damage-associated molecular patterns (DAMPs) that enhance immune cell recruitment and activation, potentially boosting ICI efficacy^[24]^. Preclinical studies demonstrate that the combination of TTFields and ICIs synergistically amplifies antitumor immunity, resulting in improved tumor control and survival outcomes^[20,25]^. Clinically, a Phase III LUNAR trial has shown that for patients with metastatic non-small cell lung cancer whose disease progresses after platinum-based chemotherapy, the addition of TTFields therapy to standard treatment (investigator’s choice of ICIs [nivolumab, pembrolizumab, or atezolizumab] or docetaxel) can enhance overall survival (OS) compared to standard treatment alone^[26]^. These findings suggest that TTFields may not only enhance ICI effectiveness but also help overcome resistance to immunotherapy in certain cancers. A clinical trial (NCT06611345) is currently underway to evaluate the efficacy and safety of combining TTFields with the TOPAZ-1 regimen for the treatment of BTC.

In our study, we applied TTFields to BTC cell lines to assess its impact on cell proliferation, migration, and mitotic processes. Notably, TTFields induced ICD, as shown by DAMP release, highlighting its potential to enhance tumor immunogenicity. Our results support the potential for TTFields to synergize with immunotherapies, offering a promising therapeutic strategy for BTC.

## RESULTS

### Validation of frequency and intensity of TTFields for human BTC cells

TTFields have been shown to be effective in killing various types of tumor cells in the frequency range of 100-300 kHz, depending on the type of tumor cell^[9,10]^. Therefore, TTFields (1.7 V/cm) at three typical frequencies of 100, 150, and 200 kHz were applied to treat two BTC cell lines, HCCC-9810 and RBE cells, for 96 h to investigate the responses of BTC cells to different frequencies. Frequency testing results showed that the inhibitory effects of TTFields were maximal at 150 kHz for these tested BTC cell lines (Fig. 1A). TTFields were applied at this optimal frequency of 150 kHz for all subsequent experiments.

**Figure 1.**
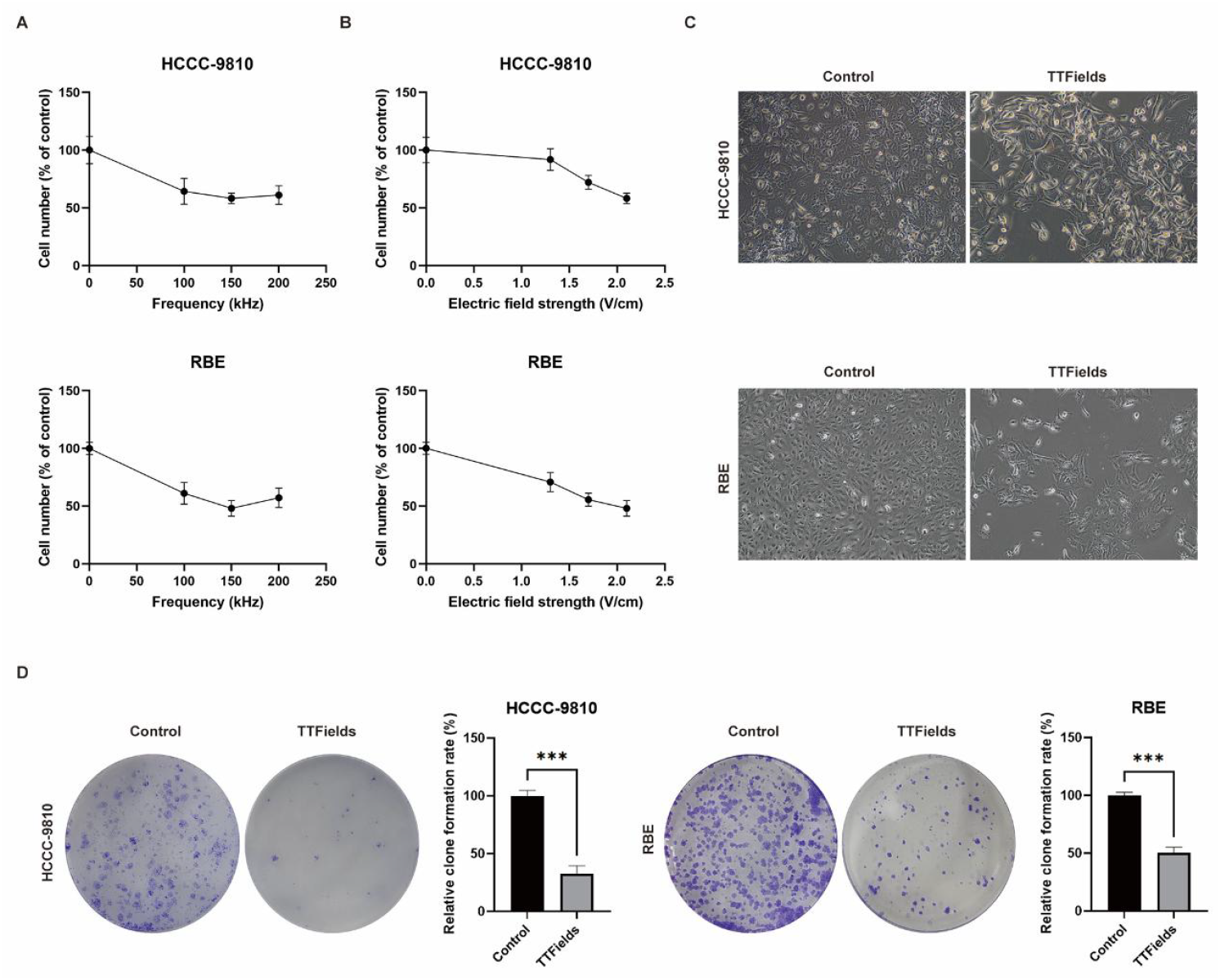
In vitro efficacy of TTFields in human BTC cells. (A) HCCC-9810 and RBE cells were treated with TTFields (1.7 V/cm) at different frequencies for 96 h and cell counts were determined. (B) Intensity-dependent effects measured after treating HCCC-9810 and RBE cells at an optimal frequency of 150 kHz for 96 h. (C) Representative images of HCCC-9810 and RBE cell morphology after 96 h of treatment with TTFields (2.1 V/cm). (D) Clonogenic potential of HCCC-9810 and RBE cell lines after treatment of TTFields (2.1 V/cm) at optimal frequency. \**p*<0.05, \*\**p*<0.01, \*\*\**p*<0.001 compared to control.

**Figure 2.**
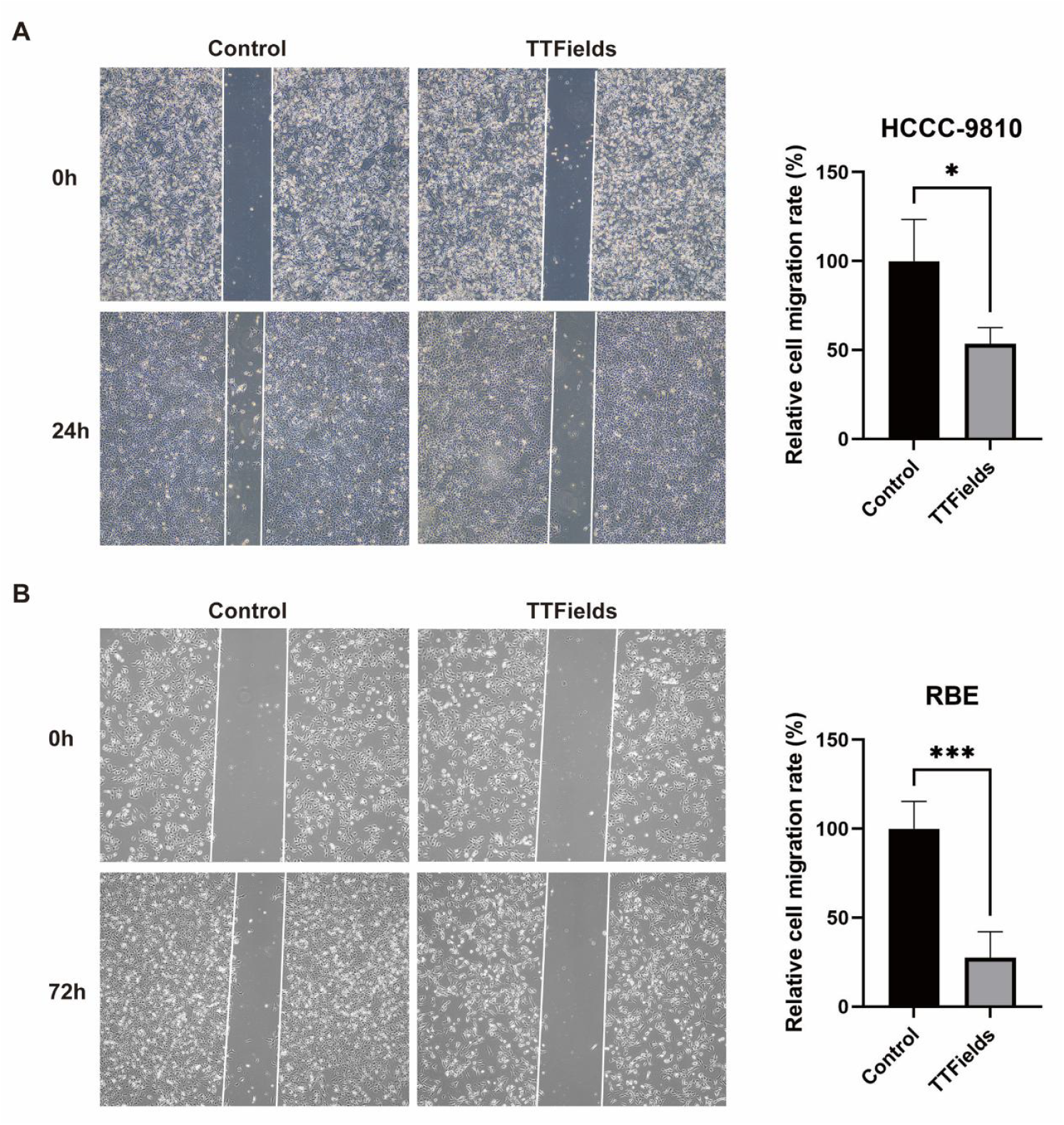
Effect of TTFields on BTC cells migration. (A) Representative images of wound healing assay of HCCC-9810 cells in different treatment groups at 0 and 24 h. (B) Quantitative data graph of wound healing assay of HCCC-9810 cells. Migration distance was calculated after 24 h of treatment with TTFields. (C) Typical images of the wound healing assay at 0 and 72 h for different treatment groups of RBE cells. (D) Plot of quantitative data of the wound healing assay of RBE cells. Migration distance was calculated after 72 h of treatment with TTFields. \**p*<0.05, \*\**p*<0.01, \*\*\**p*<0.001 compared to control.

The relationship between the intensity of TTFields at 150 kHz and cell counts was subsequently evaluated. After applying 1.3, 1.7 or 2.1 V/cm TTFields to HCCC-9810 and RBE cells for 96 h, the number of cells decreased with increasing intensity (Fig. 1B). It can be seen that the effect on cell growth depends on the intensity of TTFields. Microscopic examination revealed that the density of TTFields-treated cells was significantly reduced, and the cell morphology was changed, with larger cells and unclear edges (Figure 1C). Furthermore, the colony-forming ability of the surviving cells decreased significantly after 96 h of TTFields (2.1 V/cm) to 32.6 ± 6.8% of control for HCCC-9810 cells and to 50.4 ± 4.8% of control for RBE cells (Fig. 1D). Taken together, these results indicated that TTFields inhibited the proliferation of BTC cells.

### TTFields inhibits migration of BTC cells

Collective migration of tumor cells is the basis of cancer invasion^[27,28]^. To test the effect of TTFields treatment on the horizontal migratory capacity of BTC cells, we performed wound healing experiments. Typical wound images showed that the wound area of HCCC-9810 and RBE cells was reduced after TTFields (2.1 V/cm) treatment (Figure 3A). Quantitative data showed that the relative migration rate of HCCC-9810 cells was 53.4% after 24 h of TTFields treatment. After 72 hours of TTFields treatment, RBE cells were able to close the gap more fully, with a relative cell mobility of 27.5%. These results suggest that TTFields suppresses BTC cells migration.

**Figure 3.**
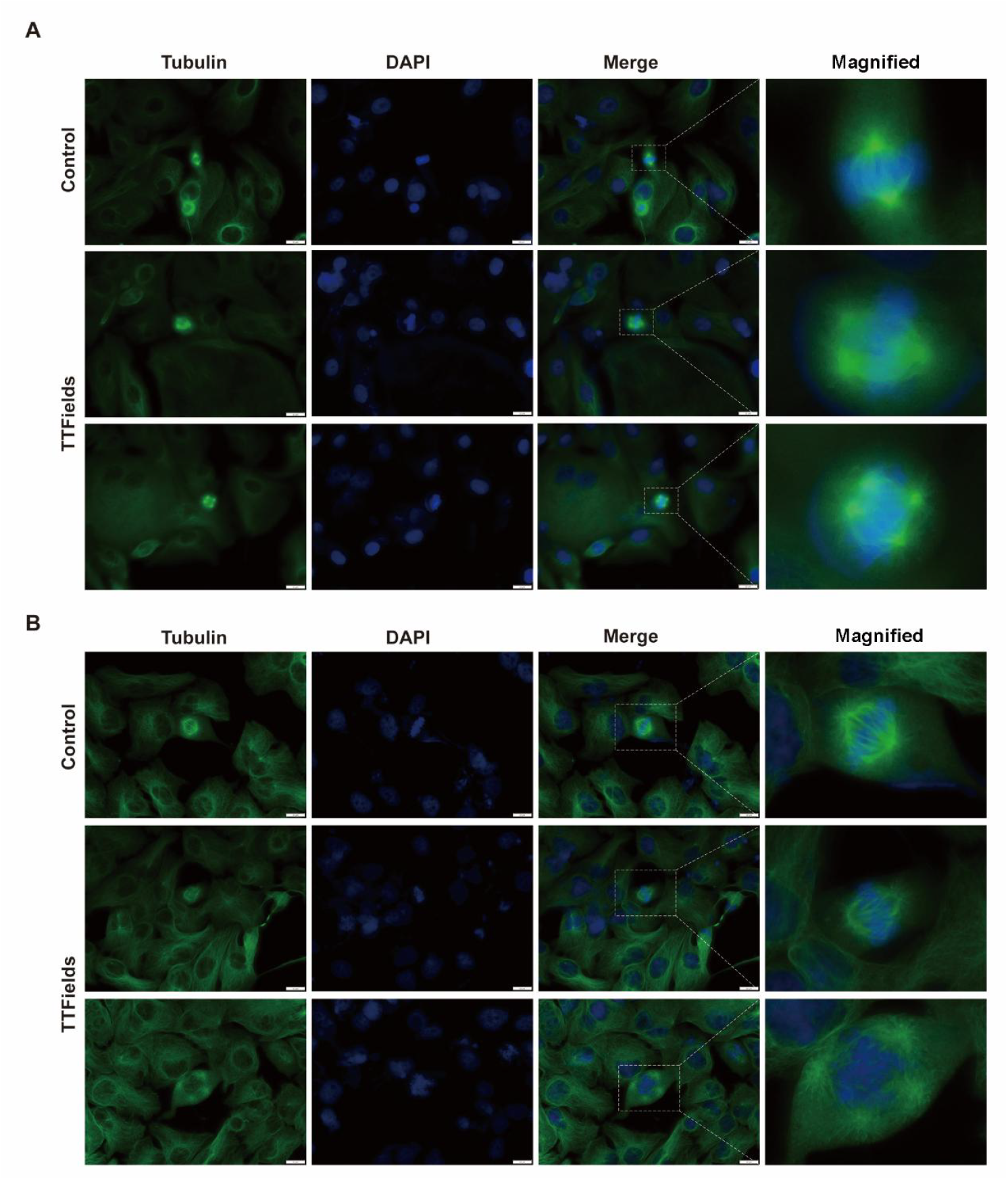
TTFields interfere with spindle morphology in mitosis of BTC cells. BTC cells were treated at 1.7 V/cm for 96 h and then stained with monoclonal antibodies for tubulin (green) and DAPI for DNA (blue), respectively. (A) Spindle morphology of HCCC-9810 cells. (B) Spindle morphology of RBE cells. Scale bars represent 20 μm.

### TTFields treatment induces aberrant mitosis in BTC cells

TTFields have been shown to disrupt the normal assembly of the mitotic spindle, thereby interfering with the replication of cancer cells^[9,29]^. To investigate whether TTFields affect normal mitosis in BTC cells, HCCC-8910 and RBE cell lines were treated with TTFields (1.7 V/cm) for 96 h. Fluorescent staining of tubulin was performed to observe changes in the spindle morphology of dividing cells. The results presented in Fig 3 indicate that both HCCC-9810 (Fig 3A) and RBE (Fig 3B) cells treated with TTFields exhibit abnormal mitosis, including disorganized spindle filament alignment (Upper panel) and multipolar mitosis (Lower panel). This suggests that TTFields treatment interferes with the normal progression of mitosis in the BTC cell line.

### TTFields treatment induces ICD in BTC cells

The extracellular release of LDH from HCCC-9810 and RBE cells increased 2-to 5-fold after 96 h of TTFields treatment. The LDH results directly indicate that TTFields treatment leads to BTC cell damage or death (Figure 4A). ICD is a mode of cell death that induces an immune response against antigens against dead-cell-related antigens, especially when they are released by cancer cells^[30-32]^. Anticancer therapies have a higher chance of success when they can also induce ICD^[33]^. It has been shown that TTFields trigger ICD in cancer cells^[25]^. To determine whether application of TTFields promotes ICD in BTC cells, HCCC-9810 and RBE cells were treated with 2.1 V/cm TTFields for 96 h and then assayed for biochemical markers of ICD that have been identified to date, including exposure of CRT at the cell membrane, extracellular secretion of ATP and HMGB1^[34]^. We first examined the surface expression of CRT after TTFields treatment using flow cytometry and observed an increase in CRT expression on the surface of BTC cells in the TTFields treatment group (Figure 4B). Next, we analyzed the release of HMGB1, another important marker of ICD. After 96 h of TTFields treatment, we found an increase in HMGB1 levels in cell supernatants (Figure 4C). In addition, extracellular ATP levels were also increased in BTC cells treated with TTFields (Figure 4D). These results suggest that TTFields may induce ICD in BTC cell lines.

**Figure 4.**
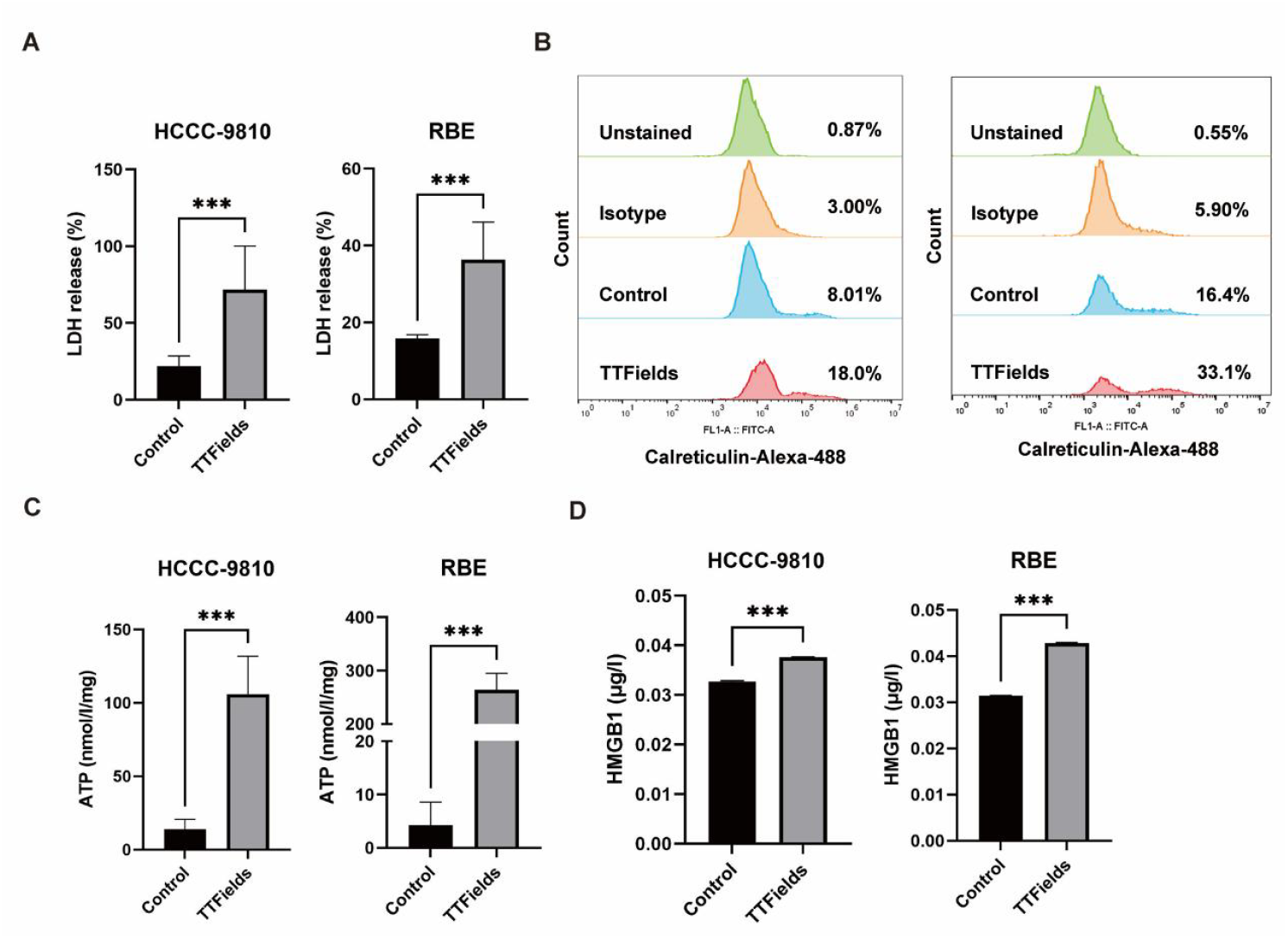
TTFields application promotes ICD in BTC cells. Conditioned media from cells treated with control or 2.1 V/cm TTFields for 96 h were collected and assayed for biochemical markers of ICD. (A) Determination of LDH release in BTC cells after TTFields treatment. (B) Membrane exposure of CRT on BTC cells was detected by flow cytometry. (C) Soluble HMGB1 in the culture medium of BTC cells was detected by ELISA kit. (D) Extracellular ATP in the culture medium of BTC cells was detected by fluorometric assay. \**p*<0.05, \*\**p*<0.01, \*\*\**p*<0.001 compared to control.

## DISCUSSION

In this study, we present the first evidence demonstrating that TTFields inhibit the proliferation of BTC cells and trigger immunogenic cell death. We identified that the frequency of 150 kHz is optimal for cytotoxicity when TTFields are applied to the HCCC-9810 and RBE human BTC cell lines, and confirmed that the efficacy of TTFields is dependent on the treatment intensity. Treatment of BTC cells with TTFields not only exhibited cytotoxic effects but also suppressed cell cloning and migration. Moreover, TTFields induced ICD in BTC cells, as shown by the heightened expression of DAMPs, such as CRT on the cell surface, as well as increased levels of HMGB1 and ATP within the conditioned medium.

TTFields exert direct effects on mitotic division in BTC cells through their alternating electric fields, as illustrated in Figure 3. The aberrant mitotic division induced by TTFields disrupts normal cell cycle progression, thereby inhibiting cellular proliferation^[35]^. This disruption is largely attributed to TTFields’ ability to interfere with microtubule polymerization, a crucial process in the formation of the mitotic spindle, which consequently obstructs the proper segregation of chromosomes during mitosis^[9,36]^. In addition to disrupting the internal structures of cancer cells, TTFields also modulate the dynamics of the cytoskeleton, which is vital for cellular motility. The changes in cytoskeletal, combined with the direct effect on the actin-myosin apparatus, impair the cancer cells’ ability to adhere and migrate efficiently^[13,37]^. As a result, TTFields significantly diminish the migratory potential of BTC cells, which is a characteristic of aggressive tumor behavior.

TTFields may induce ICD in cancer cells by inducing endoplasmic reticulum (ER) stress and initiating the unfolded protein response (UPR)^[24,38]^. This response triggers the release of DAMPs, which promote the phagocytosis of dying cancer cells by dendritic cells and macrophages. These immune cells then process and present tumor antigens to T cells, initiating an adaptive immune response^[25,30]^. TTFields-induced ICD generates an abundance of tumor antigens that promote the recruitment and activation of immune effector cells in the tumor microenvironment. When combined with immunotherapies, this synergistic approach can leverage the immunogenic properties of TTFields to jumpstart the immune system, thereby enhancing the efficacy of immunotherapeutic agents, such as ICIs, resulting in more sustained responses and prolonged survival^[39]^.

The combination of TTFields with ICIs such as anti-PD-1 or anti-PD-L1 antibodies has been explored in several preclinical models and clinical trials. For example, a separate study involving a mouse lung tumor model revealed that the application of TTFields in conjunction with ICIs significantly diminished tumor size. Notably, in groups receiving TTFields alongside anti-PD-1/anti-CTLA-4 or anti-PD-L1, there was a pronounced increase in immune cell infiltration, particularly cytotoxic T cells, into the tumor microenvironment^[20]^. Additionally, a recent study by Nitta et al. revealed that the concurrent administration of anti-PD-1 and TTFields in a GBM mouse model elicited robust myeloid and T-cell-mediated anti-tumor responses, leading to improved survival rates. This result suggests that TTFields treatment may facilitate the transformation of GBM into an “immunologically hot” tumor, thereby making it a more appealing target for immunotherapeutic approaches^[40]^. In the groundbreaking 2-THE-TOP clinical trial conducted by Novocure, the combined adjuvant treatment of temozolomide, TTFields and pembrolizumab has demonstrated promising results in patients with newly diagnosed glioblastoma. This innovative triplet therapy significantly improved PFS and OS compared to historical controls, highlighting the potential of integrating TTFields with ICIs^[41]^. The success of the TOPAZ-1 trial has catalyzed a new wave of clinical investigations exploring the potential synergistic effects between TTFields and various ICIs in BTC.

We are currently conducting a clinical trial (NCT06611345) that combines TTFields with TOPAZ-1 for the treatment of bile duct cancer. This represents the first registered clinical trial investigating TTFields in BTC. The trial will also evaluate whether TTFields enhance the efficacy of PD-1/PD-L1 inhibitors, potentially addressing major challenges in immunotherapy regarding response durability and resistance development. This research is part of the preclinical study for a clinical trial combining TTFields and TOPAZ-1 for treatment of BTC. Our findings suggest that TTFields enhance ICD in BTC cells, providing a theoretical foundation for the combination of TTFields with ICIs in BTC therapy. While these prospects are promising, it is crucial to recognize that one limitation of our study is the lack of a BTC mouse model, which hinders our capacity to thoroughly evaluate the in vivo effects of TTFields on the tumor microenvironment and immune responses. Future studies will be directed towards developing BTC-specific animal models to validate these in vitro findings and to assess the long-term efficacy of combined therapeutic approaches involving TTFields and ICIs for BTC.

In conclusion, the integration of TTFields with ICIs represents a promising therapeutic approach with potential to enhance treatment efficacy and overcome limitations of current monotherapies. With the accumulating evidence from ongoing clinical trials, it could redefine the treatment standards for a series of tumor diseases, offering more effective and personalized treatment plans tailored to cancer patients.

## MATERIALS AND METHODS

### Cell culture

Human intrahepatic cholangiocarcinoma cell lines HCCC-9810 and RBE were purchased from the Type Culture Collection of the Chinese Academy of Sciences. The cells were cultured in RPMI-1640 (Gibco, Grand Island, NY, USA) supplemented with 1% penicillin/streptomycin (pen/strep) (Gibco) and 10% heat-inactivated fetal bovine serum (FBS) (Gibco). All cell lines were maintained in a humidified chamber containing 5% CO_2_ at 37°C.

### TTFields application in vitro

HCCC-9810 cells (3 × 10^4^) or RBE cells (2 × 10^4^) were seeded on 20 mm cell culture slides and cultured for 24 or 72 h, and then the slides were transferred to 100 mm cell culture dishes (one slides/dish) using sterile forceps. The cells on the slides were exposed to an electric field strength of 1-3 V/cm, 100-250 kHz generated by a TTFields generator (Healthy Life Innovation Medical Technology, Jangsu, China). In brief, two pairs of insulated electrodes consisting of high dielectric constant ceramic sheets (ε> 3000) were inserted vertically into the cell culture dish. The electrodes were connected to sinusoidal electric potential waveform generator generating fields of the desired frequencies in the medium. The orientation of the TTFields were switched approximately 90° every 1 s. The transmission of the electric field generates a non-negligible amount of heat inside the Petri dish, dependent on the intensity of the applied field. The device with the petri dish attached is placed in an incubator and the temperature inside the petri dish is maintained at 36.5-37.5°C throughout the experiment by adjusting the temperature of the incubator. The temperature was measured by 4 thermistors attached to the ceramic walls. At the end of the treatment, the number of cells was determined using the Scepter 3.0 (Merck Millipore, Billerica, Massachusetts) automated cell counter. The relative number of cells at the end of treatment was expressed as percentage of untreated control.

### Colony formation assay

TTFields-treated HCCC-9810 and RBE cells (3000 cells/well) were inoculated into 6-well plates (WHB, Shanghai, China) for overnight growth, followed by change of cell culture medium every two days. After 7-14 days, formed colonies (cluster of 50 cells or more) were fixed with 4% formaldehyde (Solarbio, Beijing, China) throughout 30 min incubation at room temperature and then were stained with 1% Crystal Violet Solution (Beyotime Biotechnology, Shanghai, China) for 10 min. To eliminate the background staining the cells were washed twice with ddH_2_O. Colonies were photographed then counted with the Image J software.

### Wound-healing assay

The BTC cells (1.5 × 105) were seeded in cell culture slides, and subsequently grown to 75–80% confluency. A sterile 200 µl pipette tip was then used make a wound in the cells, washed the detached cells with phosphate buffer solution (PBS; Gibco), and the medium was replaced with serum-free medium. The attached cells were further incubated with TTFields at 2.1 V/cm for 24 or 72 h. Cell migration into the wound area was observed and photographed using an electron microscope (Olympus, Tokyo, Japan) at 0, 24 or 72 h. The cell migration rate was calculated using the following formula: Cell migration rate (%) = (Original width -Width after migration) × 100/Original width. Relative cell migration rate (%) = Treatment group cell migration rate/Control group cell migration rate.

### Immunofluorescence

For spindle structure analysis, HCCC-9810 and RBE cells were grown on glass culture slides and treated using the TTF system for 96 h. Cells were fixed with 4% paraformaldehyde (PFA; Solarbio) for 15 min, permeabilized in PBS (Gibco) containing 0.1% Triton X-100 and blocked with 1% bovine serum albumin (BSA; Beyotime Biotechnology) in PBS for 1 h at room temperature. The cells were stained with rabbit anti-human α-tubulin antibodies (1:800; Abcam; Cambridge, UK) overnight at 4°C. Next day, cells were then incubated with the secondary antibodies Alexa Fluor 488-conjugated anti-rabbit antibody (1:1000; Abcam; Cambridge, UK) at room temperature for 1h in the dark. Nuclei was stained with the dye 4′,6-diamidino-2-phenylindole (DAPI) (Solarbio) at 0.2 μg/ml for 10 min. Fluorescent images were captured with an Olympus IX73 fluorescence microscope (Olympus, Tokyo, Japan)

### Lactate dehydrogenase (LDH) assay

LDH activity was measured using the LDH Cytotoxicity Assay Kit following the manufacturer protocol (Beyotime Biotechnology). Briefly, all cells were incubated at 5% CO_2_, 90% humidity and 37°C for 24 h. Subsequently, 120 µl supernatant per well was carefully transferred into the corresponding wells of a clear 96-well plate. Add a total of 60 µl of reaction mixture to each well and incubate for 30 min at room temperature (∼25°C) away from light. Absorbance was measured at 490 nm on a Tecan Spark® 20M (Tecan, Mannedorf, Switzerland). Cytotoxicity (LDH release rate) was calculated according to the following formula: Cytotoxicity (%) = (test sample - blank control) × 100/(maximum enzyme activity control - blank control).

### Calreticulin cell surface expression assays

HCCC-9810 and RBE cells were collected and stained with rabbit polyclonal anti-calreticulin antibody (1:100; Abcam; Cambridge, UK) or isotype control rabbit IgG (1:100; Abcam; Cambridge, UK) in flow cytometry buffer (1% BSA in PBS) for 45 min. Cells were then washed and incubated with anti-rabbit Alexa Fluor 488 conjugated antibody (1:2000; Abcam; Cambridge, UK) in flow cytometry buffer for 30 min. Labeled cells were subsequently detected using a BD Accuri C6 flow cytometer (BD Biosciences, San Jose, CA, USA). Data were analyzed using FlowJo Software version 10 (TreeStar, San Francisco, CA, USA).

### ATP and HMGB1 assays

Supernatants from HCCC-9810 and RBE cells treated with TTFields for 96 h were collected for HMGB1 and ATP release assays. ATP concentrations in cell supernatants were measured by an enhanced ATP assay kit (Beyotime Biotechnology) according to the manufacturer’s instructions. The HMGB1 levels in the supernatant of treated cells were analyzed by an enzyme-linked immunosorbent assay (ELISA) kit (Meimian, Shanghai, China) according to the manufacturer’s protocol. Luminescence and absorbance were measured using a Tecan Spark® 20M microplate multimode reader (Tecan, Mannedorf, Switzerland).

## Statistical Analysis

Dates are presented as the mean ± standard deviation (SD). The statistical analyses and figures were made by GraphPad Prism version 9 (GraphPad Software, Inc., San Diego, CA, USA). Statistical significance was defined by unpaired two-tailed Student’s *t*-tests. The analysis of multiple groups was performed with ANOVA with an appropriate post hoc test. *p* < 0.05 was considered statistically significant and indicated as \**p* < 0.05; \*\**p* < 0.01; and \*\*\**p* < 0.001.

## AUTHOR CONTRIBUTIONS

Y.S., M.Y. and Y.Y. conceived and designed the study. Y.Y. conducted the experiments and collected the data. Y.Y. and Y.W. performed data analysis and interpretation. Y.S. and Y.Y. wrote the initial manuscript draft. All authors critically reviewed and approved the final version of the manuscript for submission.

## FUNDING

This study is supported by Healthy Life Innovation Medical Technology Co., Ltd.

## CONFLICT OF INTEREST

Y.Y., Y.W. and Y.S are employees of Healthy Life Innovation Medical Technology Co., Ltd, a startup developing medical devices for treating tumors. The authors declare that this affiliation does not influence the design, execution, or interpretation of the research presented in this study.

## ACKNOWLEDGMENTS

Authors thank Dr. Min Yao, Dr. Mingwu Deng, Ms. Meixin Han, Ms. Huihui Huang, Ms. Jinye Yang, Mr. Jiajie Hui, and Mr. Sheng Chen from Healthy Life Innovation Medical Technology Co., Ltd for assistance on experiments and technical support on experimental devices.

## REFERENCE

[1] Valle, J. W., Kelley, R. K., Nervi, B.Oh, D.-Y. & Zhu, A. X. Biliary tract cancer. Lancet 397, 428–444 (2021).

[2] Nakamura, H. et al. Genomic spectra of biliary tract cancer. Nat Genet 47, 1003–1010 (2015).

[3] Valle, J. W., Lamarca, A., Goyal, L., Barriuso, J. & Zhu, A. X. New Horizons for Precision Medicine in Biliary Tract Cancers. Cancer Discov 7, 943–962 (2017).

[4] Lamarca, A., Edeline, J. & Goyal, L. How I treat biliary tract cancer. ESMO Open 7, 100378 (2022).

[5] Oneda, E., Abu Hilal, M. & Zaniboni, A. Biliary Tract Cancer: Current Medical Treatment Strategies. Cancers (Basel) 12, 1237 (2020).

[6] Leowattana, W., Leowattana, T. & Leowattana, P. Paradigm shift of chemotherapy and systemic treatment for biliary tract cancer. World J. Gastrointest. Oncol 15, 959–972 (2023).

[7] Kang, S., El-Rayes, B. F. & Akce, M. Evolving Role of Immunotherapy in Advanced Biliary Tract Cancers. Cancers (Basel) 114, 1748 (2022).

[8] Oh, D.-Y. et al. Durvalumab or placebo plus gemcitabine and cisplatin in participants with advanced biliary tract cancer (TOPAZ-1): updated overall survival from a randomised phase 3 study. Lancet Gastroenterol Hepatol 9, 694–704 (2024).

[9] Kirson, E. D. et al. Disruption of cancer cell replication by alternating electric fields. Cancer Res 64, 3288–3295 (2004).

[10] Kirson, E. D. et al. Alternating electric fields arrest cell proliferation in animal tumor models and human brain tumors. Proc Natl Acad Sci U S A 104, 10152–10157 (2007).

[11] Wenger, C. et al. A Review on Tumor-Treating Fields (TTFields): Clinical Implications Inferred From Computational Modeling. IEEE Rev Biomed Eng 11, 195–207 (2018).

[12] Moser, J. C. et al. The Mechanisms of Action of Tumor Treating Fields. Cancer Res 82, 3650–3658 (2022).

[13] Voloshin, T. et al. Tumor Treating Fields (TTFields) Hinder Cancer Cell Motility through Regulation of Microtubule and Acting Dynamics. Cancers (Basel) 12, 3016 (2020).

[14] Mehta, M., Wen, P., Nishikawa, R., Reardon, D. & Peters, K. Critical review of the addition of tumor treating fields (TTFields) to the existing standard of care for newly diagnosed glioblastoma patients. Crit Rev Oncol Hematol 111, 60–65 (2017).

[15] Xing, Y. et al. Advancements and current trends in tumor treating fields: a scientometric analysis. Int J Surg 110, 2978–2991 (2024).

[16] Seufferlein, T., Gracian, A. C. & Kueng, M. PANOVA-4 (EF-39): Pilot study of tumor treating fields (TTFields) therapy with atezolizumab, gemcitabine (GEM), and nab-paclitaxel (NabP) as first-line (1L) treatment for metastatic pancreatic ductal adenocarcinoma (mPDAC). JCO 42, TPS718 (2024).

[17] Ceresoli, G. L., Vergote, I., Rivera, F. & Pless, M. Abstract CT202: Safety of Tumor Treating Fields delivery to the torso: Pooled analysis from TTFields clinical trials. Cancer Res 79, CT202–CT202 (2019).

[18] Mehta, M. P. et al. Results from METIS (EF-25), an international, multicenter phase III randomized study evaluating the efficacy and safety of tumor treating fields (TTFields) therapy in NSCLC patients with brain metastases. JCO 42, 2008 (2024).

[19] Davidi, S. et al. Tumor Treating Fields (TTFields) Concomitant with Sorafenib Inhibit Hepatocellular Carcinoma In Vitro and In Vivo. Cancers (Basel) 14, 2959 (2022).

[20] Barsheshet, Y. et al. Tumor Treating Fields (TTFields) Concomitant with Immune Checkpoint Inhibitors Are Therapeutically Effective in Non-Small Cell Lung Cancer (NSCLC) In Vivo Model. Int J Mol Sci 23, 14073 (2022).

[21] Yi, M. et al. Combination strategies with PD-1/PD-L1 blockade: current advances and future directions. Mol Cancer 21, 28 (2022).

[22] Yue, S., Zhang, Y. & Zhang, W. Recent Advances in Immunotherapy for Advanced Biliary Tract Cancer. Curr Treat Options Oncol 25, 1089–1111 (2024).

[23] Casak, S. J. et al. FDA Approval Summary: Durvalumab and Pembrolizumab, Immune Checkpoint Inhibitors for the Treatment of Biliary Tract Cancer. Clin Cancer Res 30, 3371–3377 (2024).

[24] Lan, Y., Zhang, S., Pan, Y., Wang, M. & Chen, G. Research Progress on the Mechanism of Anti-Tumor Immune Response Induced by TTFields. Cancers (Basel) 15, 5642 (2023).

[25] Voloshin, T. et al. Tumor-treating fields (TTFields) induce immunogenic cell death resulting in enhanced antitumor efficacy when combined with anti-PD-1 therapy. Cancer Immunol Immunother 69, 1191–1204 (2020).

[26] Leal, T. et al. Tumor Treating Fields therapy with standard systemic therapy versus standard systemic therapy alone in metastatic non-small-cell lung cancer following progression on or after platinum-based therapy (LUNAR): a randomised, open-label, pivotal phase 3 study. Lancet Oncol 24, 1002–1017 (2023).

[27] Friedl, P. & Gilmour, D. Collective cell migration in morphogenesis, regeneration and cancer. Nat Rev Mol Cell Biol 10, 445–457 (2009).

[28] Trepat, X., Chen, Z. & Jacobson, K. Cell migration. Compr Physiol 2, 2369–2392 (2012).

[29] Giladi, M. et al. Mitotic Spindle Disruption by Alternating Electric Fields Leads to Improper Chromosome Segregation and Mitotic Catastrophe in Cancer Cells. Sci Rep 5, 18046 (2015).

[30] Kroemer, G., Galassi, C., Zitvogel, L. & Galluzzi, L. Immunogenic cell stress and death. Nat Immunol 23, 487–500 (2022).

[31] Gao, X. et al. Disulfiram/Copper Induces Immunogenic Cell Death and Enhances CD47 Blockade in Hepatocellular Carcinoma. Cancers (Basel) 14, 4715 (2022).

[32] Decraene, B. et al. Immunogenic cell death and its therapeutic or prognostic potential in high-grade glioma. Genes Immun 23, 1–11 (2022).

[33] Turubanova, V. D. et al. Immunogenic cell death induced by a new photodynamic therapy based on photosens and photodithazine. J Immunother Cancer 7, 350 (2019).

[34] Kroemer, G., Galluzzi, L., Kepp, O. & Zitvogel, L. Immunogenic cell death in cancer therapy. Annu Rev Immunol 31, 51–72 (2013).

[35] Giladi, M. et al. Mitotic disruption and reduced clonogenicity of pancreatic cancer cells in vitro and in vivo by tumor treating fields. Pancreatology 14, 54–63 (2014).

[36] Gera, N. et al. Tumor treating fields perturb the localization of septins and cause aberrant mitotic exit. PLoS One 10, e0125269 (2015).

[37] Xiao, T., Zheng, H., Zu, K., Yue, Y. & Wang, Y. Tumor-treating fields in cancer therapy: advances of cellular and molecular mechanisms. Clin Transl Oncol 27, 1–14 (2025).

[38] Shteingauz, A. et al. AMPK-dependent autophagy upregulation serves as a survival mechanism in response to Tumor Treating Fields (TTFields). Cell Death Dis 9, 1074 (2018).

[39] Han, Y., Tian, X., Zhai, J. & Zhang, Z. Clinical application of immunogenic cell death inducers in cancer immunotherapy: turning cold tumors hot. Front Cell Dev Biol 12, 1363121 (2024).

[40] Nitta, R. T., Lee, S. Y., Lim, M. & Li, G. Tumor treating fields (TTFields) can induce immunogenic cell death in GBM resulting in enhanced immune modulation. Cancer Res 84, 7513–7513 (2024).

[41] Tran, D. D., Ghiaseddin, A., Chen, D. & Le, S. B. Final analysis of 2-THE-TOP: A phase 2 study of TTFields (Optune) plus pembrolizumab plus maintenance temozolomide (TMZ) in patients with newly diagnosed glioblastoma. JCO 41, 2024 (2023).

